# 3D Human Neuromuscular Architecture Encoded by Compliant Patterned Hydrogel Scaffolds

**DOI:** 10.64898/2026.09.25.754309

**Authors:** Asia Badolato, Aiste Balciunaite, Elena Casalino, Ilakkiya Ravindrarajah, Miriam Filippi, Melanie Generali, Thomas Laumonier, Robert K. Katzschmann

## Abstract

Engineered human neuromuscular models are limited by a trade-off between biological fidelity and architectural control: self-assembled organoids organize stochastically, while engineered 3D tissues impose anisotropy through passive tension at the edges without internal guidance. Here we encode architectural guidance into human neuromuscular tissues by coupling a three-dimensional, cell-laden biomatrix to a xolographically printed hydrogel substrate featuring sub-millimeter surface topography. We demonstrate that grooved substrates increase human primary myotube alignment and maturation in the overlying 3D tissue, while guiding directional axon extension from human iPSC-derived motor neuron spheroids. We show that combining both cell types along a shared alignment axis establishes functional neuromuscular connectivity, evidenced by acetylcholine receptor clustering, glutamate-evoked contraction and its blockade by d-tubocurarine (curare), and yields innervation coverage of centimeter scale tissues. Overall, this platform combines compliant substrates, mesoscale tissue architectural guidance, and human-relevant cell biology in a single untethered construct for studies of neuromuscular tissue development and function.

## 1. Introduction

Neuromuscular diseases collectively affect millions of people worldwide and remain without curative treatment.^[1–4]^ A central challenge to therapy development is the lack of experimental models that faithfully reproduce the structural and functional organization of native human neuromuscular tissue. Skeletal muscle is hierarchically organized and strongly anisotropic: myofibers are grouped into parallel fascicles that are mechanically coupled through the extracellular matrix,^[5–7]^ and are innervated by motor axons that run longitudinally alongside the fibers within intramuscular nerve branches before terminating at neuromuscular junctions (NMJs).^[8–12]^ This spatial organization underlies efficient motor unit recruitment, graded force generation and stable neuromuscular transmission,^[13,14]^ and its disruption drives progressive functional decline in pathological states. Understanding how tissue architecture governs neuromuscular function therefore requires experimental models that not only employ appropriate human cell sources but also encode the architectural cues guiding neuromuscular tissue formation and maintenance in vivo.

Stem cell engineering has enabled human neuromuscular organoids, assembloids and compartmentalized co-cultures that model diseases or report drug responses.^[15–21]^ Despite their biological fidelity, self-assembled systems depend on stochastic cellular organization, which limits spatial control over tissue architecture and reproducibility of neuromuscular geometry.^[22,23]^ In contrast, engineered platforms that couple cell-laden hydrogels to flexible posts or microfabricated frames impose mechanical constraints that promote muscle alignment, and this alignment can in turn provide directional cues for axon extension along the tissue’s longitudinal axis, indicating that tissue-scale anisotropy can itself guide motor axon growth.^[24–33]^ However, these tethered architectures anchor the tissue to posts or supporting structures, constraining the muscle’s contractile behavior and creating mechanical mismatch between the tissue and its supporting scaffold. These limitations indicate that combining tissue-level anisotropy with spatially uniform innervation requires guidance cues to be distributed throughout the tissue architecture, rather than imposed solely through mechanical tethering at the tissue periphery.

This motivates a distinction between two mechanistically different topographic cues: contact and architectural guidance. Contact guidance acts locally through micron or sub-micron scale surface topography at the cell-substrate interface,^[34–39]^ whereas architectural guidance operates at the mesoscale to organize myofibers and axons throughout a three-dimensional tissue. The first therefore acts at the cell level, in direct contact with the patterned surface, while the second can establish anisotropic organization across a volumetric muscle tissue, guiding myofiber alignment and axonal extension as the tissue forms and remodels. Extending topographic guidance from the cell-substrate interface to the full three-dimensional tissue volume at the centimeter scale therefore represents a largely unexplored opportunity for engineering biomimetic NMJ models, enabling architectural cues to guide neuromuscular organization throughout the tissue. Realizing this approach with human cells to generate human NMJ models would address an important gap in the field, with implications for both the biological fidelity and fabrication scalability of existing human NMJ models.

Here we present a human neuromuscular platform in which the mesoscale surface topography of a soft substrate encodes architectural guidance into a three-dimensional, cell-laden biomatrix throughout tissue formation and maturation. Using xolography as the volumetric printing method,^[40,41]^ we fabricate compliant hydrogel substrates mechanically matched to native skeletal muscle stiffness and patterned with parallel sub-millimeter grooves at the centimeter length scale.^[42]^ We first show that groove topography independently enhances 3D myofiber maturation and organization and promotes guided axon bundle extension from motor neuron spheroids. We then show that combining both cell types on a grooved substrate in a parallel configuration establishes functional NMJ connectivity, validated by acetylcholine receptor clustering, neuron-evoked contraction and pharmacological blockade with d-tubocurarine. Finally, by comparing parallel and orthogonal muscle-neuron geometries, we show that placing the neuronal sources at both ends of the fiber axis increases innervation coverage across centimeter scale constructs. Collectively, this platform establishes a structured human neuromuscular tissue model that combines architectural guidance, 3D matrix mechanics and human cell biology, providing a versatile foundation for investigating NMJ formation and neuromuscular disease, and for future applications in biohybrid actuation.

## 2. Results and Discussion

We previously established a co-optimized bilayer volumetric system in which sub-millimeter grooved hydrogel scaffolds are coupled with an engineered murine muscle layer through a continuous interface. ^[42]^ Here we adapt this platform for human neuromuscular models, incorporating dedicated seeding compartments for each cell type and leveraging surface topography to instruct both muscle and neural maturation along a shared alignment axis to generate functional neuromuscular tissues.

### 2.1 Volumetrically printed grooved substrates stably couple to 3D human neuromuscular tissue

Micro architected scaffolds were fabricated using xolography, a volumetric dual-color printing method combining high geometrical freedom, rapid fabrication and micrometer resolution,^[40]^ using a previously reported formulation of polyethylene glycol diacrylate (PEGDA) 700 and gelatine methacrylate (GelMA).^[42]^ The design was adapted to contain a central region patterned with parallel grooves (300 µm wide, 500 µm deep) with rectangular profile,^[42]^ and two chambers at the groove ends used as motor neuron seeding sites and designed to align axon bundles along the muscle fiber direction (**Figure 1A**). We confirmed that surface topography remained stable under culture-like conditions by measuring groove width and depth over 7 days in PBS at 37 °C (**Figure S1A**). Scaffold-conditioned medium was applied to motor neuron cultures to assess potential cytotoxicity of fabrication byproducts, confirming that the scaffold affected neither cell viability nor metabolic activity (**Figure S1B**).

**Figure 1.**
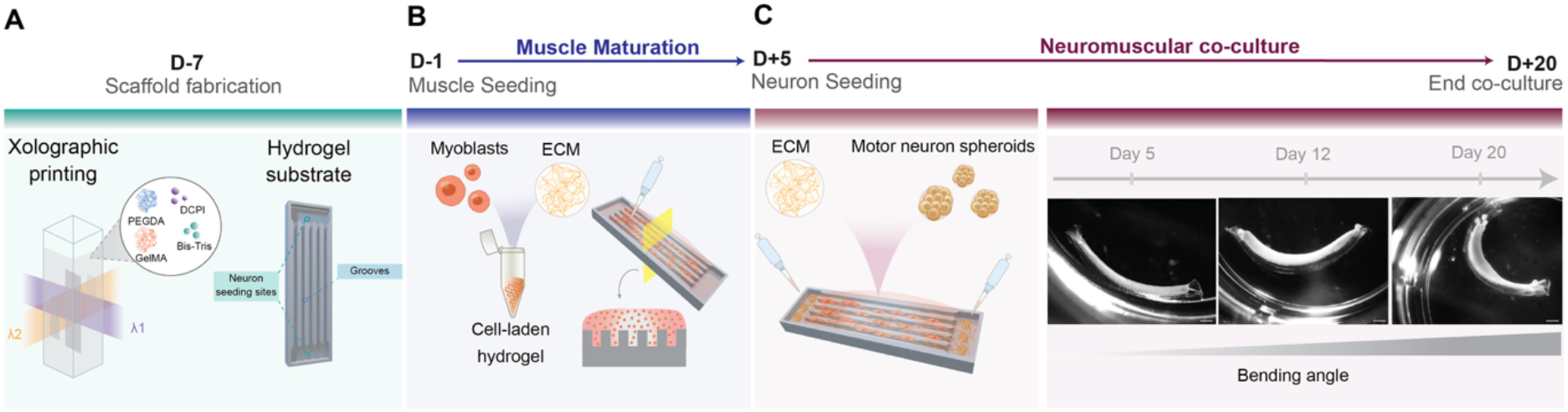
Fabrication process of hydrogel substrates coupled to a 3D human neuromuscular tissue. **(A)** Scaffold fabrication using dual-wavelength light-sheet printing of a PEGDA-GelMA precursor containing Bis-Tris as co-initiator and a dual-color photoinitiator (DCPI). The scaffold design features a central region of parallel grooves and two motor neuron seeding sites at the groove ends. **(B)** Human primary myoblasts suspended in a fibrin-Matrigel precursor are deposited onto the grooved region and crosslinked in situ, forming a 3D cell-laden layer that fills and spans the grooves. Constructs are maintained in muscle differentiation medium for 5 days. **(C)** Left: pre-differentiated human iPSC-derived motor neuron spheroids are placed in the dedicated compartments at both ends of the groove axis and embedded in the same matrix. The co-culture is maintained until day +20. Right: stereomicroscopy of a construct at days 0, 7 and 15 of co-culture, showing progressive bending of the substrate as the tissue compacts and remodels. Scale bars, 1 mm.

To couple a 3D engineered tissue with the scaffold, an extracellular matrix (ECM)-like hydrogel based on fibrin and Matrigel and loaded with myoblasts was deposited and crosslinked on the scaffold’s surface (**Figure 1B**). While cell monolayers adhere and spread readily on PEGDA-GelMA through its native adhesion motifs, a 3D tissue additionally requires stable anchoring of the hydrogel matrix to the substrate. We therefore functionalized the scaffolds with a polydopamine (PDA) coating before seeding, a strategy shown to promote adhesion of fibrin and collagen hydrogels to inert polymer surfaces.^[43–45]^ Coated substrates retained the hydrogel after crosslinking, whereas uncoated surfaces delaminated upon medium addition (**Figure S2**). After 5 days of tissue maturation, pre-differentiated motor neuron spheroids were added to the dedicated compartments of the scaffold, covered with a layer of fibrin-Matrigel. The resulting constructs were maintained in co-culture medium for an additional 14 days **(Figure 1C)**. As the tissue developed and remodeled, tissue-generated forces induced scaffold bending. The compliant scaffold supported this deformation while maintaining intact tissue-scaffold interfaces.

To characterize the mechanical properties underlying this behavior, we performed rheological measurements, which showed that the printed scaffold had a storage modulus approximately one order of magnitude higher than that of the fibrin-Matrigel matrix (∼1.5 kPa vs. ∼200 Pa; **Figure S3A**). Thus, the scaffold combines sufficient stiffness to mechanically support the developing tissue with a compliant architecture that allows bending in response to passive forces generated during tissue remodeling and contraction. Under tensile loading, the scaffold exhibited a Young’s modulus of ∼25kPa (**Figure S3B**), within the reported range for transversal modulus of skeletal muscle ^[46,47]^,indicating that its tensile mechanical properties fall within the range found in vivo. Importantly, assuming incompressibility, G’ ≈ 1.5 kPa corresponds to E ≈ 4.5 kPa, so we hypothesize that the remaining difference reflects the distinct specimen geometries and measuring methods applied. To assess whether this mechanical design was compatible with functional tissue development, the constructs were electrically stimulated at the end of the maturation period. Upon stimulation, the constructs exhibited global bending in response to muscle contraction and recovered their original shape following relaxation (**Video S1**). Having established that this material system supports functional tissue development, we next investigated the effects of surface topography on muscle and neural tissues separately.

### 2.2. Mesoscale grooves align and mature myofibers throughout the 3D tissue layer

To assess how surface topography influences myofiber organization within the volumetric muscle layer, we compared tissues cultured on grooved and flat scaffolds. After 14 days, the tissue-scaffold interface was preserved under both conditions, independently of surface topography (**Figure 2A, B**). On grooved substrates, however, myotubes formed elongated, unidirectionally aligned bundles running parallel to the grooves, in contrast to the more disorganized network on flat substrates (**Figure 2C**). Quantitative analysis of myotube alignment confirmed a significantly greater myotube orientation along the groove direction, whereas myotubes on flat substrates exhibited broadly distributed alignment angles (**Figure 2D**). To establish that this organization is not restricted to the groove walls, we measured myotube orientation as a function of distance from the nearest groove wall. Alignment was uniform across the full 150 µm half-width, from the wall to the groove center, and exceeded the flat-substrate control at every distance (**Figure 2E**). The consistent alignment across the groove width indicates that the influence of the topographical cue extends beyond the immediate vicinity of the groove walls. Grooved substrates further showed higher fusion index and greater fiber density (**Figure 2F, G**), consistent with enhanced myofiber formation and more uniform tissue development. Together, these results demonstrate that the topographical cues are transmitted to the overlying 3D tissue layer and promote directional organization with associated improvements in structural features of myogenic maturation. We next investigated whether the same scaffold topography could similarly guide motor neuron growth and axon bundle extension.

**Figure 2.**
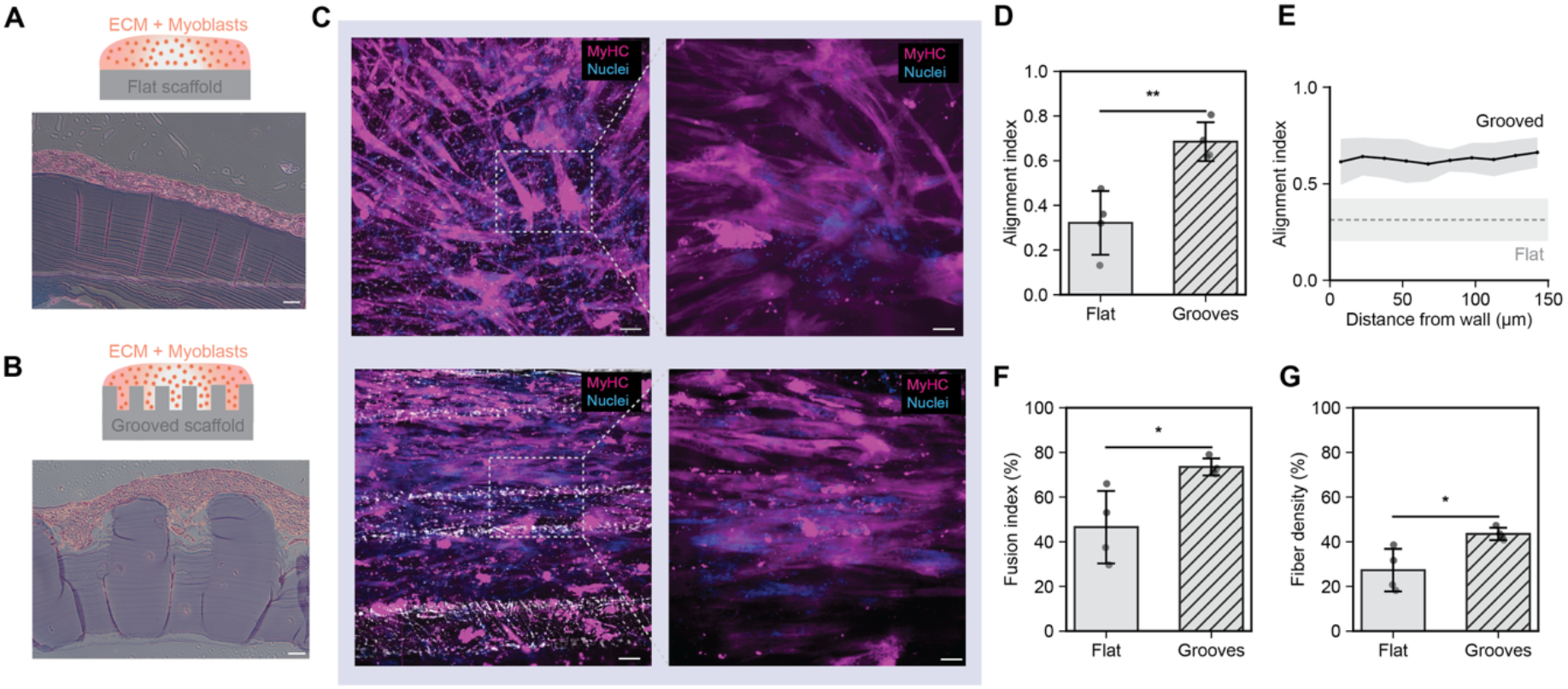
Mesoscale grooves align and mature myofibers throughout the 3D tissue layer. **(A, B)** Schematics and hematoxylin-and-eosin-stained cross-sections of muscle tissue on flat (A) and grooved (B) scaffolds after 14 days of culture, showing a continuous tissue-scaffold interface in both conditions and tissue filling the groove cavities. Scale bars, 100 µm. **(C)** Maximum-intensity projections of myosin heavy chain (MyHC, magenta) and nuclei (blue) for tissues on flat (top) and grooved (bottom) substrates. Right panels show magnifications of the boxed regions. Scale bars, 100 µm (overviews) and 50 µm (insets). **(D)** Myotube alignment index relative to the groove axis (grooved) or the corresponding reference axis (flat). **(E)** Myotube alignment as a function of distance from the nearest groove wall, across the full 150 µm groove half-width. **(F)** Fusion index of myoblasts after a total of 20 days of culture on flat and grooved substrates. **(G)** Fiber density as percentage of the field of view area after a total of 20 days of culture on flat and grooved substrates. Data are represented as mean ± SD. Each point represents one independent biological replicate. Statistical significance was calculated using unpaired two-tailed Welch’s t-test; *p < 0.05, **p < 0.01.

### 2.3. Groove topography directs and promotes axon outgrowth from motor neuron spheroids

In vivo, motor axons are organized into densely packed fascicles that support long-range signal transmission and muscle innervation.^[48–50]^ We therefore asked whether the scaffold could support motor neuron culture while promoting directional axon growth. Human iPSC-derived motor neuron spheroids were matured in suspension for 7 days and then cultured for an additional 15 days on either standard 2D glass substrates or on scaffolds (**Figure 3A**). On 2D glass substrates, spheroids extended axons isotropically and expressed key neuronal maturation markers including βIII-tubulin, ChAT and HB9, after 22 days of total culture (**Figure 3B**). When embedded in a fibrin-Matrigel matrix on the scaffold, spheroids retained expression of these markers while extending axons preferentially along the groove direction, suggesting that the topographical cues promote directional axon growth without apparent impairment of neuronal maturation. Moreover, glutamate stimulation elicited a rapid increase in intracellular calcium in motor neuron spheroids cultured on both glass and scaffolds, indicating that the neurons retained a functional response to excitatory stimulation (**Figure S4, Video S2**).

**Figure 3.**
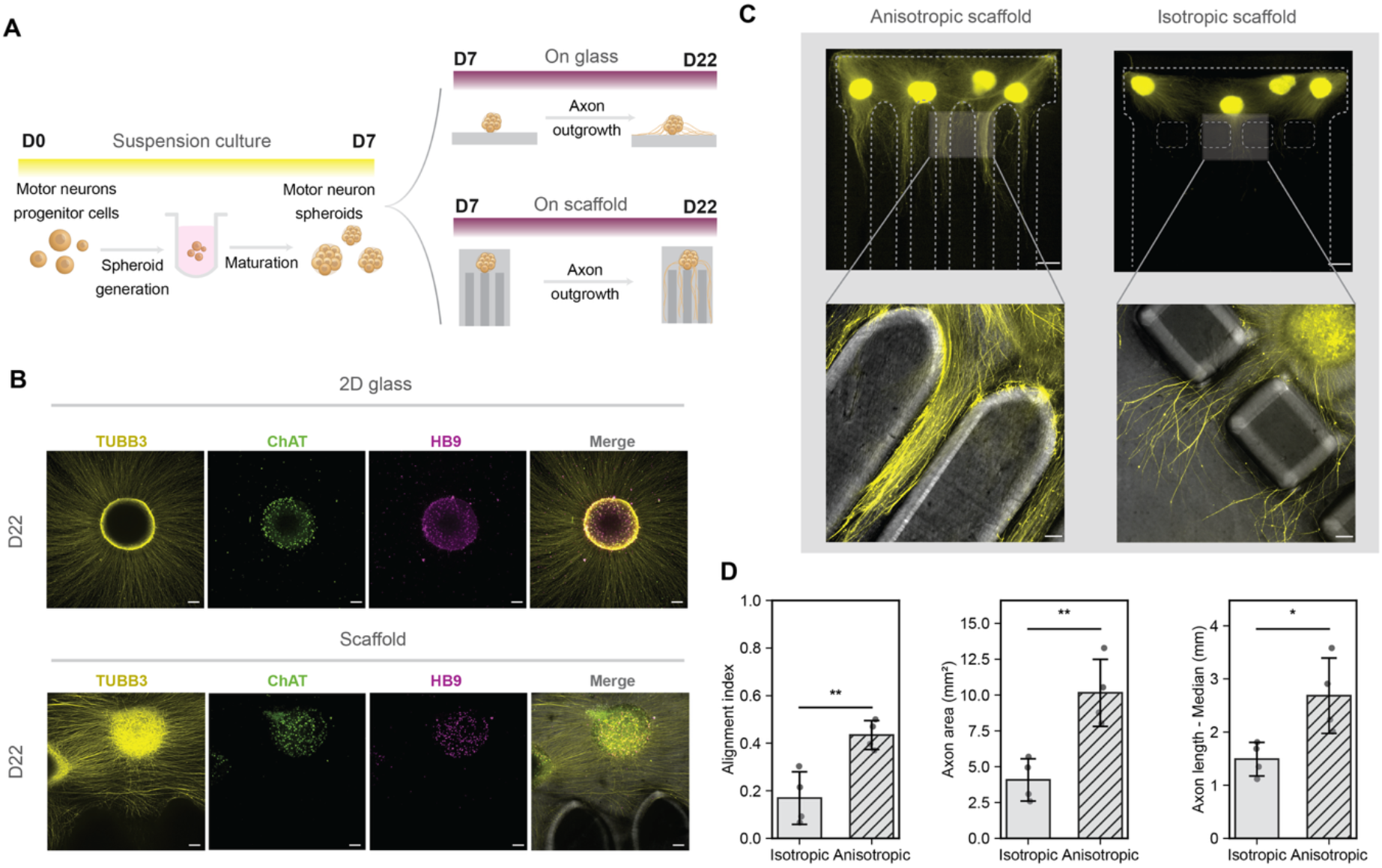
Groove topography directs and promotes axon outgrowth from motor neuron spheroids. **(A)** Experimental timeline: human iPSC-derived motor neuron progenitors are aggregated in spheroids and matured in suspension for 7 days, then cultured for a further 15 days either on 2D glass or embedded in fibrin-Matrigel on xolographically printed scaffolds. **(B)** Immunostaining at day 22 for βIII-tubulin (TUBB3, yellow), choline acetyltransferase (ChAT, green) and HB9 (magenta) for spheroids on 2D glass (top) and on scaffolds (bottom). Scale bars, 100 µm. **(C)** Top: stereomicroscopy of TUBB3 signal from spheroids seeded on anisotropic (grooved, left) and isotropic (right) scaffolds with equal seeding area. Bottom: confocal magnifications of the boxed regions overlaid on brightfield. Scale bars, 0.5 mm (overviews) and 100 µm (insets). **(D)** Axon alignment index (left), axon-covered area (center) and median axon length (right) on isotropic and anisotropic scaffolds. Data are represented as mean ± SD. Each point represents one independent biological replicate. Statistical significance was calculated using unpaired two-tailed Welch’s t-test; *p < 0.05, **p < 0.01.

To isolate the contribution of groove geometry to axon extension, we compared anisotropic and isotropic scaffold geometries while maintaining the same spheroid-seeding area. In anisotropic scaffolds, axon bundles extended from the spheroid core along the parallel grooves, whereas on isotropic scaffolds axons were initially channeled out of the seeding area and then dispersed without preferential directionality (**Figure 3C**). Quantitative analysis confirmed a higher alignment index in the anisotropic condition (**Figure 3D**), demonstrating that groove topography promotes directional axon extension resembling native nerve organization. Axon-covered area and axon length were also increased in the presence of grooves (**Figure 3D**), indicating that the surface topography influences not only axon alignment but also axon extension. These features may be advantageous for guiding axonal outgrowth toward and across centimeter scale muscle constructs.

Having established that the grooved architecture independently supports muscle organization and motor neuron axon extension, we next asked whether combining both cell types on the same scaffold could promote functional neuromuscular coupling.

### 2.4. Architected scaffolds support functional neuromuscular junction formation

To generate a neuromuscular co-culture, myoblasts were seeded onto the scaffold as described above and differentiated in monoculture. After 5 days of differentiation, pre-matured motor neuron spheroids were seeded into the designated compartments with additional hydrogel matrix to promote stable adhesion, and the co-cultures were maintained for a total of 15 days before analysis (**Figure 4A**). As a first structural indicator of NMJ formation, we quantified acetylcholine receptor (AChR) clustering on muscle fibers, a hallmark of postsynaptic specialization, in constructs culture with and without motor neurons (**Figure 4B**). Co-cultures containing motor neurons (+MN) showed significantly more AChR clusters than muscle-only controls (−MN) (**Figure 4C**), indicating enhanced postsynaptic organization in the presence of motor neurons. To assess whether these structural features were associated with functional neuromuscular connectivity, we measured muscle calcium activity before and after neuron-specific glutamate stimulation. Muscle-only cultures showed no significant change in the number of spontaneously active regions after glutamate addition, whereas co-cultures exhibited a marked increase (**Figure 4D**), consistent with transmission of neuronal excitation to the muscle through functional neuromuscular connections. Accordingly, localized contractile regions adjacent to motor neuron spheroids contracted synchronously following glutamate addition, consistent with synaptic activation (**Figure S5, Video S3**). Beyond these local responses, glutamate stimulation also evoked synchronous contractions that propagated across macroscopic regions of the tissue (**Figure 4E, Video S4**). These contractions were blocked by addition of curare, a competitive antagonist of nicotinic acetylcholine receptors (**Figure 4F**), supporting the involvement of cholinergic neuromuscular transmission in muscle activation and demonstrating functional neuromuscular coupling at the tissue scale.

**Figure 4.**
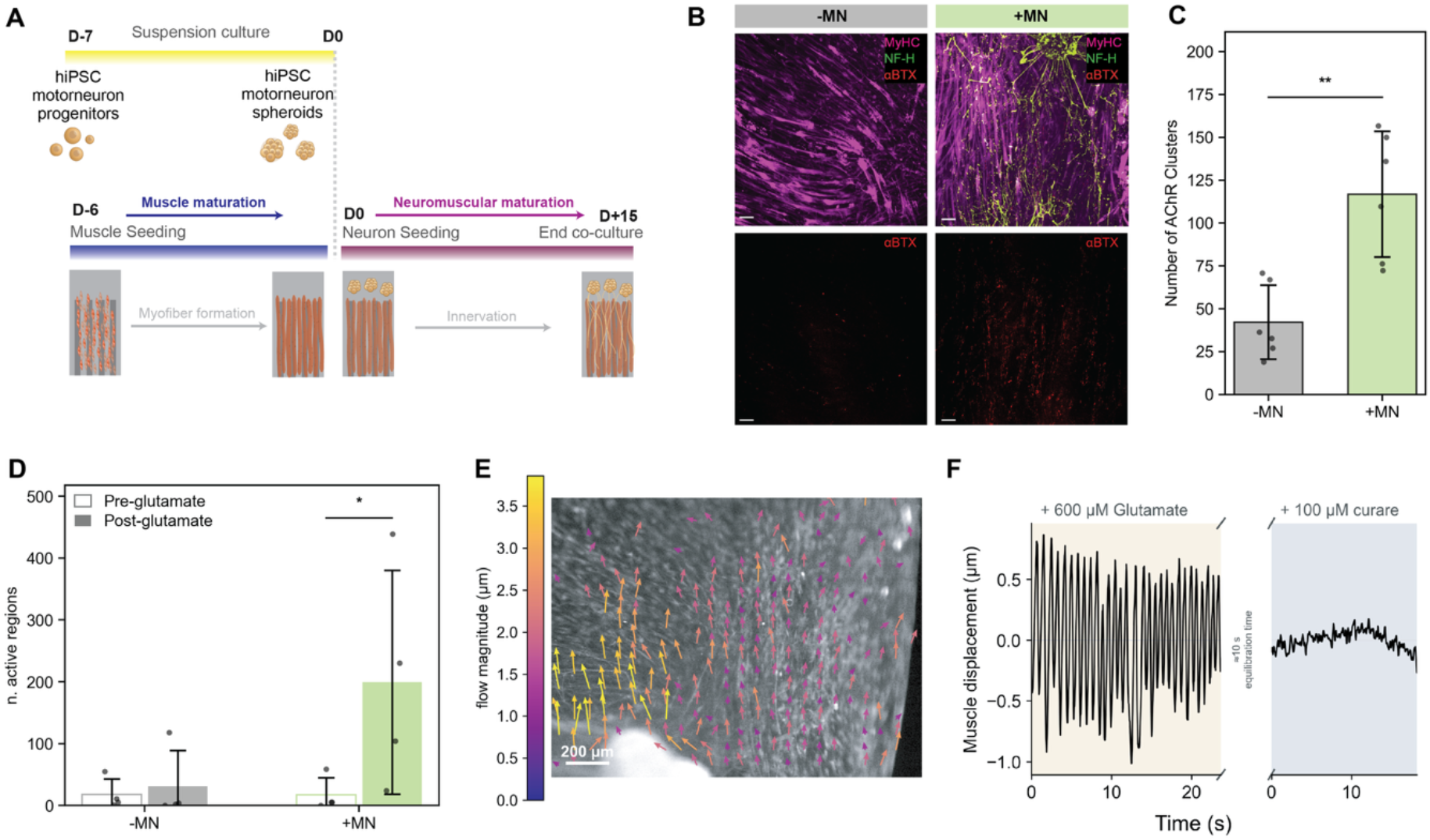
Architected scaffolds support functional neuromuscular junction formation. **(A)** Co-culture timeline: motor neuron spheroids are matured in suspension from day −7 to day 0 while myoblasts seeded at day −6 are matured in monoculture. Spheroids are then added at day 0 and the co-culture is maintained for 15 days. **(B)** Myosin heavy chain (MyHC, magenta), neurofilament heavy chain (NF-H, green) and α-bungarotoxin (αBTX, red) staining of muscle-only (−MN) and co-cultured (+MN) constructs; bottom, αBTX channel alone. Scale bars, 100 µm. **(C)** Number of AChR clusters averaged per field of view in −MN and +MN constructs. **(D)** Number of spontaneously active calcium regions before (pre-glutamate) and after (post-glutamate) addition of 600 µM L-glutamate, for −MN and +MN constructs. **(E)** Optical-flow field of a co-culture after glutamate addition, showing coordinated displacement propagating across the construct. Arrow color and length encode flow magnitude. Scale bar, 200 µm. **(F)** Representative muscle displacement trace after addition of 600 µM glutamate (left) and after subsequent addition of 100 µM curare (right), showing inhibition of the evoked contraction. Data are represented as mean ± SD. Each point represents one independent biological replicate. Statistical significance of independent samples was calculated using unpaired two-tailed Welch’s t-test; *p < 0.05, **p < 0.01. Statistical significance of paired samples (the same construct imaged before and after glutamate stimulation) was calculated using a two-tailed paired t-test on log-transformed values, with *p < 0.05, **p < 0.01.

### 2.5. Muscle-neuron spatial configuration and groove topography enhances innervation efficiency

To determine whether the parallel muscle-neuron configuration provides advantage for muscle innervation, we compared it with a muscle geometry in which neuron spheroids were positioned along the long edge of the muscle with axons initially extending orthogonally to the fiber direction.^[34]^ Scaffolds for both configurations were designed with equivalent muscle area and number of neuron spheroids. In both configurations, axons projected toward the muscle, but their trajectories differed: in the orthogonal configuration axons deviated upon reaching the muscle and subsequently realigned with the fiber direction, whereas in the parallel configuration axons extended along the muscle axis and remained aligned with the fibers without substantial deflection (**Figure 5A**). Innervation coverage, quantified as the ratio of axon-positive to total muscle area, was significantly greater in the parallel configuration (**Figure 5B**). Consistent with this, plotting local axon coverage together with the muscle area present at each distance (**Figure S6**) shows that, in the orthogonal configuration, coverage declined to a small fraction of its near-source value while muscle tissue was still abundant, whereas in the parallel configuration coverage persisted across the full section range. This difference may arise from the spatial distribution of the motor neuron sources. In the parallel configuration, spheroids are placed at both ends of the muscle along its long axis, providing opposing sources of axonal outgrowth and increasing the spatial extent of muscle accessible to innervation. In the orthogonal configuration, spheroids are distributed along one edge, resulting in more localized distribution of axonal input. Together, these results indicate that positioning motor neuron spheroids at opposing ends of the muscle can enhance the extent and uniformity of innervation in centimeter scale muscle constructs.

**Figure 5.**
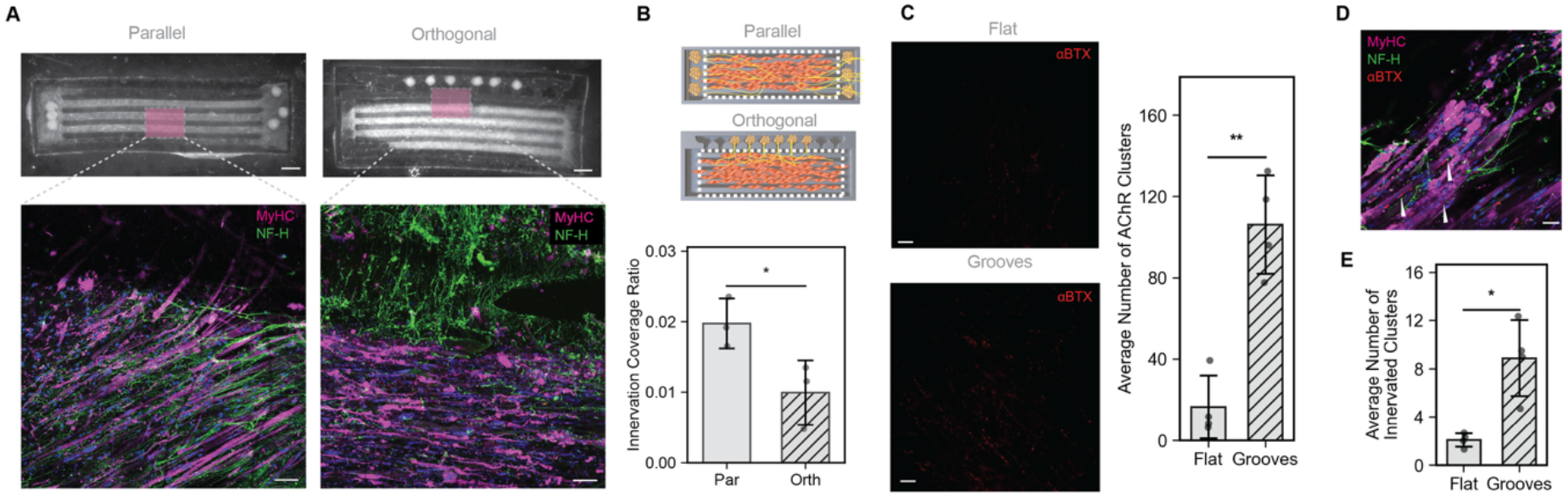
Muscle-neuron spatial configuration and groove topography determine innervation efficiency. **(A)**Stereomicroscopy overviews (top) and confocal magnifications (bottom) of constructs in parallel vs orthogonal configurations showing expression of MyHC (magenta) and NF-H (green). Scale bars, 1 mm (overviews) and 100 µm (insets). (B) Schematics of the two configurations highlighting the innervated area (top) and innervation coverage (bottom), quantified as the ratio of axon-positive regions within total muscle area. **(C**) αBTX staining of co-cultures on flat (top) and grooved (bottom) substrates, with the corresponding average number of AChR clusters. Scale bars, 100 µm. **(D)** Representative confocal image of merged MyHC (magenta), NF-H (green), αBTX (red), and nuclei (dapi) channels showing receptor clusters overlapping to motor axons (arrowheads). Scale bar, 100 µm. **(E)** Number of innervated regions calculated as co-localized αBTX/NF-H structures per field of view on flat and grooved substrates. Data are represented as mean ± SD. Each point represents one independent biological replicate. Statistical significance of independent samples was calculated using unpaired two-tailed Welch’s t-test, and using *p < 0.05, **p < 0.01.

We next examined whether substrate topography further influenced innervation in co-culture by comparing grooved and flat substrates in co-culture. Postsynaptic structures identified by αBTX-positive AChR clusters on the muscle fibers were higher on grooved substrates (**Figure 5C**), consistent with the enhanced muscle organization reported above and with previous studies showing that contact guidance on anisotropic surfaces promotes AChR clustering.^[35,51]^ To assess whether motor axons were associated with these postsynaptic clusters, we quantified co-localization of αBTX/NF-H co-localization as a readout of putative NMJ sites (**Figure 5D**), as previously reported.^[52]^ Grooved scaffolds yielded significantly more putative innervated clusters than flat controls (**Figure 5E**), confirming that scaffold topography is associated with formation of neuromuscular contact sites.

## 3. Conclusions

We present a human neuromuscular platform in which micro scale surface topography, volumetrically printed into a mechanically compliant hydrogel, encodes architectural guidance into an overlying three-dimensional tissue. Relative to flat substrates of identical composition and stiffness, grooved scaffolds increased human myotube alignment and maturation and promoted axon bundle elongation from human iPSC-derived motor neuron spheroids. Combined in co-culture along this shared axis, the two cell types formed acetylcholine receptor clusters and contracted in response to glutamate, a response subsequently blocked by curare, indicating functional neuromuscular coupling. The constructs also contracted globally under bulk electrical stimulation while remaining free of posts or frames, yielding a centimeter scale innervated muscle in a single untethered format. Critically, this guidance was not confined to the cell-substrate interface: myotube alignment was uniform from the groove wall to the groove center and exceeded the flat control at every distance. We therefore propose that grooves act as mechanical boundary conditions on the hydrogel volume rather than as a contact cue: by constraining how the cell-laden matrix compacts and transmits tension during remodeling, they establish an anisotropic stress field that orients fibers throughout the layer, with axon bundles following the same pathways and showing corresponding gains in alignment and extension. Topographical patterning of a soft substrate can therefore organize an entire tissue volume, while providing a compliant tissue-scaffold interface. We further identified the spatial configuration of muscle and neurons as a design parameter for innervation. Axons entering from one long edge deviated on contact with the muscle and realigned with the fibers, whereas spheroids placed at both ends of the long axis projected axons that ran continuously along the fibers and increased innervation coverage with the same number of spheroids in a more compact design. The platform thus separates and places under design control two variables that are typically coupled in engineered neuromuscular tissue: the mechanical boundary conditions imposed on the tissue and the geometry of its innervation, both encoded in a single printing process that accommodates a wide range of geometrical variations. The result is an untethered human neuromuscular platform at centimeter scale, in which architecture is a design variable, laying the foundation for modelling neuromuscular disease in patient-derived cells and for biohybrid actuators controlled by neural inputs.

## 4. Experimental Section

### 4.1. Human primary myoblast culture

Human primary myoblasts were isolated from human skeletal muscle biopsies as previously described. ^[53]^ Muscle biopsies were obtained under a protocol approved by the Commission Cantonale d’Éthique de la Recherche of the Canton of Geneva, Switzerland (project “Myogenic stem cells and improvement of muscle regeneration,” approval no. PB_2016-01793, PI: Dr. T Laumonier; approved on 18 October 2018). Written informed consent was obtained from all participants in accordance with the guidelines and regulations of the Swiss health authorities. Following isolation, human myoblasts were expanded in growth medium (GM): DMEM/F-12 supplemented with 20% FBS, 1% penicillin-streptomycin (PS), dexamethasone (0.4 µg mL^−1^), human epidermal growth factor (hEGF, 10 ng mL^−1^), basic fibroblast growth factor (bFGF, 1 ng mL^−1^) and human insulin (10 µg mL^−1^). Monoculture of human myoblasts were differentiated in differentiation medium (DM1) consisted of high-glucose DMEM supplemented with 1% PS, 6-aminocaproic acid (ACA, 1 mg mL^−1^), BSA (0.5 mg mL^−1^), uridine (50 µg mL^−1^), creatine (1 mM) and insulin (10 µg mL^−1^).

### 4.2. Motor neuron culture

Human iPSC-derived motor neurons (3020M1-1M, Anatomic Incorporated) were thawed and seeded at 5000 cells per well in ultra-low-attachment 96-well plates, then centrifuged to promote aggregation. Spheroids were matured in suspension in Moto-MM medium (Anatomic) for 1 week according to the manufacturer’s instructions, then transferred to adherent culture on ibidi µ-Slides or seeded directly onto scaffolds. During adherent culture, motor neurons were maintained in co-culture maturation medium (hDM2): 1:1 Neurobasal: DMEM/F-12 supplemented with 1% PS, ACA (1 mg mL^−1^), 1× B27, 1× N2, 1× GlutaMAX, uridine (50 µg mL^−1^), creatine (1 mM), insulin (5 µg mL^−1^), ascorbic acid (50 µg mL^−1^), and BDNF, GDNF and CNTF (10 ng mL^−1^ each).

### 4.3. Scaffold fabrication

Scaffolds were designed in Fusion 360 (Autodesk) and printed by xolography on a Xube^2^ volumetric printer (xolo GmbH, Germany). The hydrogel precursor consisted of PEGDA (Mn 700), GelMA (Rousselot X-Pure 160P80), Bis-Tris (Sigma-Aldrich, ≥99.0%, #B9754) and dual-color photoinitiator (DCPI 5004, xolo GmbH) at final concentrations of 15 v/v%, 5 wt%, 500 mM and 0.3 mg mL^−1^, respectively. Printing cuvettes (Brand UV 759170) were filled with precursor and gelled at 16 °C for 30 min before printing at the same temperature.

After printing, cuvettes were warmed to 37 °C to dissolve uncured resin and scaffolds were isolated in PBS, rinsed extensively and incubated at 37 °C for 1 h to dissolve residual uncured material. Scaffolds were then incubated in 0.1 wt% lithium phenyl-2,4,6-trimethylbenzoylphosphinate (LAP) for 30 min and post-cured under a 405 nm lamp for 15 min. Scaffolds were rinsed in PBS and stored at 4 °C for up to one week before seeding.

### 4.4. PDA coating and scaffold post-processing

Dopamine hydrochloride was dissolved at 1 mg mL^−1^ in 10 mM Tris-HCl buffer (pH 8.5). Scaffolds were incubated in this solution for 1 h at 37 °C to allow polydopamine deposition, after which the solution was aspirated, and the scaffolds rinsed three times in deionized water. Scaffolds were then dried under UV light in a biosafety cabinet for 1 h.

### 4.5. Mechanical characterization

Rheological characterization was performed on an Anton Paar MCR 302 rheometer (Anton Paar, Graz, Austria) with an 8 mm parallel-plate geometry. Disk specimens (8 mm diameter, 1 mm thickness) were printed and post-processed as described for scaffold fabrication, then incubated in hGM at 37 °C overnight before testing. Time sweeps were acquired at a constant frequency of 1 Hz, 0.5% strain and 25 °C over 100 points; the storage modulus was calculated as the mean of the last 80 points.

Tensile tests were performed on an MTS system with a 100 N force transducer (MTS Systems Corporation, model 661.09B-21). Dog-bone specimens with widened clamping regions were printed from the same formulation and by the same process as the scaffolds; the gauge section measured 12 mm (length) × 3.5 mm (width) × 3 mm (thickness). Samples were stored in water and tested hydrated at room temperature. Specimens were attached to the clamps with cyanoacrylate glue and mounted between the grips. Displacement was increased linearly to 12 mm over 2000 s (0.36 mm min^−1^), with a rupture detector terminating the test after failure. Force and displacement were recorded throughout. Engineering stress was calculated as force divided by the initial cross-sectional area of the gauge section, and engineering strain as displacement divided by the initial gauge length. Young’s modulus was determined from the slope of the linear region of the stress-strain curve.

### 4.6. Co-culture generation on scaffolds

Human primary myoblasts at passage 5 or 6 were detached with 0.05% trypsin-EDTA and resuspended at 15 × 10^6^ cells mL^−1^ in a hydrogel precursor of fibrinogen (4 mg mL^−1^), Matrigel (30 v/v%), thrombin (0.05 U/mg of fibrinogen) and hGM. The cell-laden hydrogel was pipetted onto the grooved compartment and polymerized at 37 °C for 1 h before hGM was added. After 1 day in hGM, the medium was switched to hDM1 and half-refreshed daily for 5 days.

After 5 days of muscle maturation, six motor neuron spheroids of 300-400 µm diameter were manually seeded per scaffold into the designated site, covered with the same fibrin-Matrigel hydrogel, and polymerized for 30 min at 37 °C. The medium was then switched to hDM2 and refreshed every other day for 15 days of co-culture.

### 4.7. Calcium imaging and chemical stimulation

Cells and tissues were loaded with Fluo-4 AM or Calbryte-520 (5 µM) in Live Cell Imaging Solution (LCIS) containing 0.04% Pluronic F-127 for 1 h at 37 °C, rinsed in LCIS and equilibrated for 30 min at room temperature. Recordings were acquired on a Zeiss Stemi 508 stereomicroscope with a GFP module at ≥20 fps. For chemical stimulation of motor neurons, L-glutamic acid was applied at a final concentration of 600 µM. For AChR blockade, curare was applied at a final concentration of 100 µM.

### 4.8. Electrical stimulation

For global electrical stimulation, constructs were placed in a custom plate fitted with fixed graphite electrodes connected to a function generator and submerged in high-glucose DMEM. Constructs were stimulated with square-wave pulses at 1 Hz, 20 V and 0.1% duty cycle. Live recordings were acquired on a stereomicroscope at ≥20 fps.

### 4.9. Histology

After in vitro culture, the constructs were fixed in 4% paraformaldehyde (PFA) at room temperature for 20 min. After several washes in PBS, the samples were embedded in paraffin and sectioned at 4.5 µm thickness with a microtome (Microm HM430, Thermo Fisher Scientific). Sections were stained with hematoxylin (GHS116, Sigma-Aldrich) and eosin (HT110116, Sigma-Aldrich), mounted, and imaged in brightfield with a light microscope (Olympus CKX41, Olympus Schweiz AG).

### 4.10. Immunocytochemistry

After fixing, samples were permeabilized in 0.2% Triton X-100 for 30 min, blocked in 1% BSA / 1% Tween-20 for 1 h at room temperature, and incubated with primary antibodies (Table S1A) diluted in blocking buffer overnight at 4 °C. Samples were then rinsed in PBS and incubated with secondary antibodies and DAPI (Table S1B) for 3 h at room temperature. Confocal imaging was performed on a Nikon Ti2 confocal microscope.

### 4.11. Statistical analysis

Data are presented as mean ± SD unless otherwise stated, with each data point representing one independent biological replicate (averaged across fields of view). Comparisons among independent or dependent conditions were performed using an unpaired, two-tailed Welch’s *t*-test and a two-tailed paired t-test on log-transformed values, respectively. Analyses were performed in Python (SciPy, statsmodels) with significance set at *p < 0.05, **p < 0.01.

## Acknowledgements

The work was supported by the SNSF Sinergia Grant No. 216727. At ETH Zurich, this work was done within the framework of the ALIVE initiative (Advanced Engineering with Living Materials) and funded by the SFA-AM program (Strategic Focus Area — Advanced Manufacturing). The authors thank Dr. Costanza Giampietro and Prof. Dr. Edoardo Mazza for the support and use of the confocal microscope, Prof. Dr. Mark Tibbitt and his group members for the support and use of the rheometer and plate reader, and the members of the DBM Histology Core Facility (University of Basel) for support and execution of the experiments.

## Author Contributions

A.B. conceived the study, designed and performed the experiments, analyzed the data and wrote the manuscript. A.Ba. contributed to experimental design and methodology; E.C. and I.R. contributed to cell culture and mechanical characterization; M.F. and M.G. contributed to methodology and resources; T.L. provided human primary myoblasts; R.K.K. supervised the study and acquired funding. All authors reviewed and approved the final manuscript.

## Data Availability

All data supporting the findings of this study are available within the article and its Supporting Information. Raw imaging datasets and the custom Python analysis pipelines are available from the corresponding author upon reasonable request.

