## Supplementary Figures for "3D Human Neuromuscular Architecture Encoded by Compliant Patterned Hydrogel Scaffolds"

### SUPPLEMENTARY INFORMATION

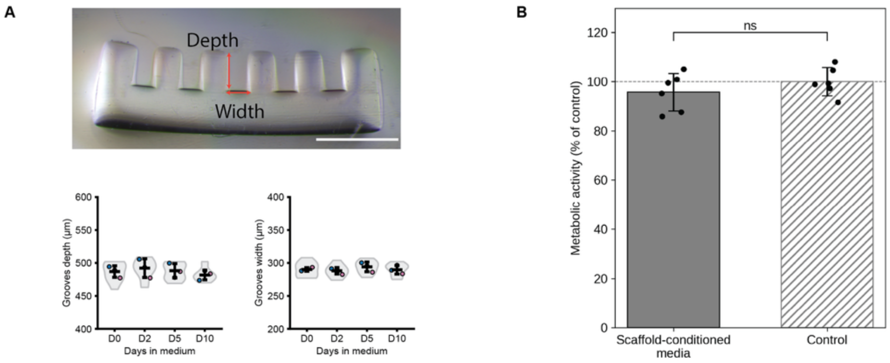


**Figure S1.** Dimensional stability and cytocompatibility of the volumetrically printed scaffold. (A) Representative stereomicroscope image of a scaffold's cross-section, indicating the groove depth and width measured (top - scale bar, 1 mm), and quantification of groove depth and width after 0, 2, 5 and 10 days of incubation in culture medium (bottom). Each point is one cross-section (n = 3); bars show mean ± SD. (B) Metabolic activity of hiPSC-derived motor neuron spheroids cultured for 48 h in scaffold-conditioned medium, normalized to cells in fresh medium (control, dashed line at 100%). Each point is one biological replicate (n = 6); bars show mean ± SD; ns, not significant (Welch's test).

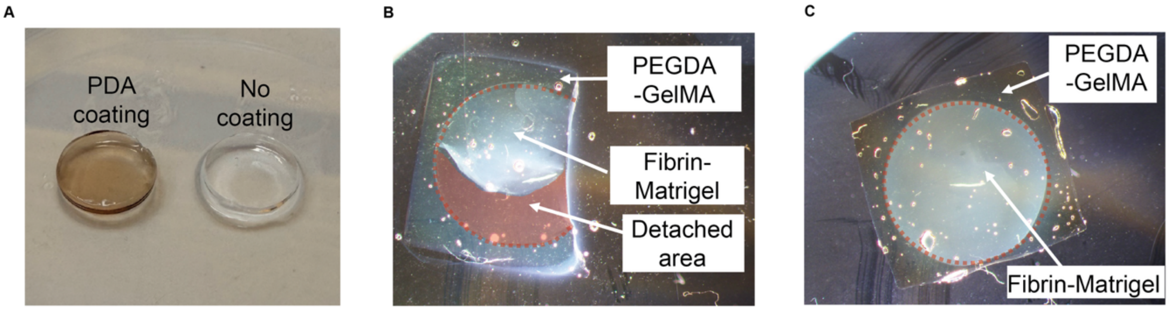


**Figure S2.** Polydopamine coating is required for stable adhesion of the fibrin-Matrigel layer to the printed scaffold. (A) Representative images of PEGDA-GelMA hydrogels after polydopamine (PDA) treatment (left, characteristic brown color) and without coating (right). (B) Uncoated PEGDA-GelMA molds were casted with fibrin-Matrigel hydrogel and placed at 37°C for 1 h to allow crosslinking. Upon medium addition, the casted layer retracts immediately from the scaffold surface indicating low attachment. The dashed outline marks the originally cast area and the shaded region the detached area. (C) PDA-coated models processed in parallel: the fibrin–Matrigel layer remains fully adherent across the same area over the entire culture period.

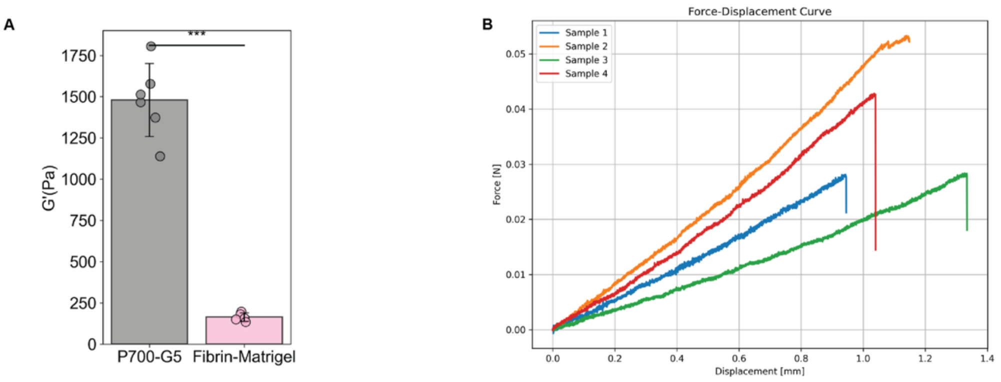


**Figure S3.** Mechanical characterization of the printed scaffold and of the cell-laden matrix. (A) Storage modulus (G') of printed P700-G5 and fibrin-Matrigel samples. Each data point is one independent sample (n = 6) bars show mean ± SD; ***$p < 0.001$ (unpaired two-tailed t-test). (B) Force-displacement curves from uniaxial tensile testing of four independent scaffolds, where drop in each curve marks macroscopic failure.

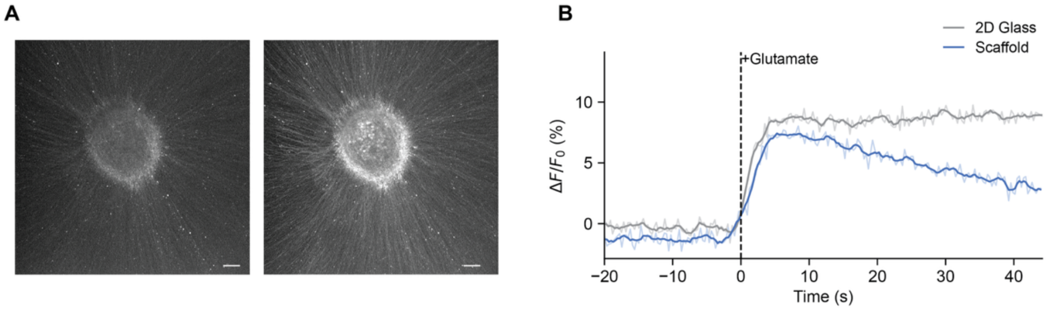


**Figure S4.** Motor neuron spheroids response to glutamate. (A) Representative images of calcium response from spheroids on glass before (left) and after (right) glutamate stimulation Scale bars, 100 µm. (B) Quantification of calcium response to glutamate stimulation (added at t=0, dashed line), expressed as $\Delta F/F_0$ of the signal averaged over the spheroid.

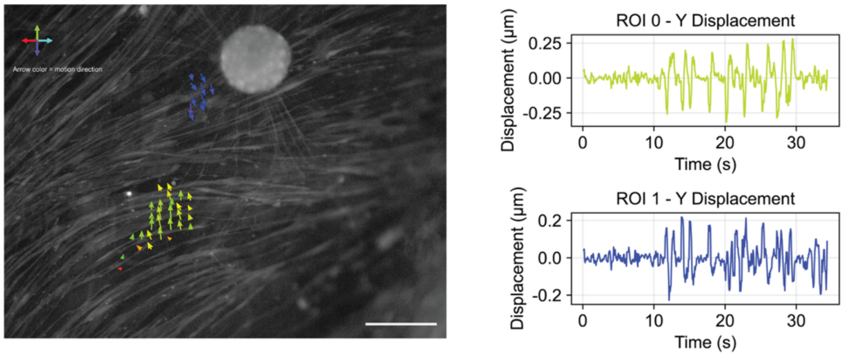


**Figure S5.** Contractile local activity. Left: representative frame of the muscle with optical-flow displacement vectors overlaid in two regions of interest. Arrow color encodes the direction of motion. Scale bar, 300 µm. Right: y-displacement traces extracted from the two regions of interest over 30 s of recording, showing repetitive twitches of sub-micrometer amplitude occurring synchronously in both regions.

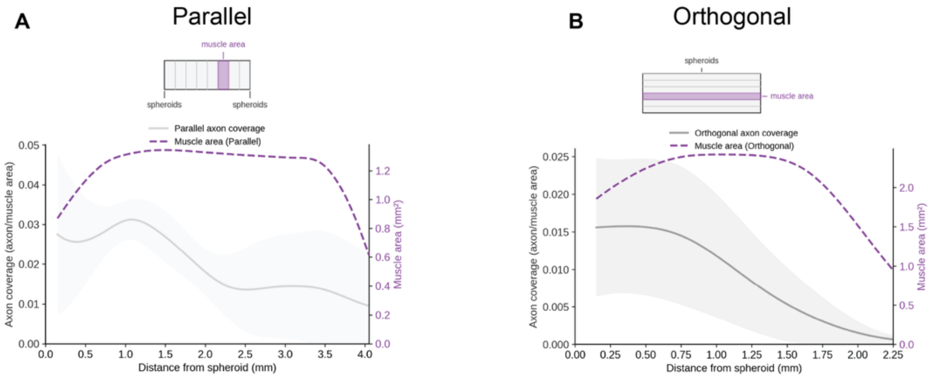


**Figure S6.** Axon coverage falls before muscle tissue's width in the orthogonal configuration. Local axon coverage plotted together with the muscle area in that section (purple dashed, right axis), as a function of distance from the nearest spheroid, for the parallel (A) and orthogonal (B) configurations. The schematic above each panel defines a section for that geometry: sections across the width of the construct for the parallel configuration, and sections along its length for the orthogonal configuration. Curves are means of n = 3 constructs per condition, smoothed for display; shaded bands show ± SD. The orthogonal profile ends at 2.25 mm because no muscle tissue extends beyond the width of the construct.
